# Hydrogel microdroplet based glioblastoma drug screening platform

**DOI:** 10.1101/2025.07.08.663758

**Authors:** Brittany A. Payan, Annika Carrillo Diaz De Leon, Tejasvi Anand, Gunnar B. Thompson, Ana Mora-Boza, Vishnu V. Krishnamurthy, Andrés J. García, Brendan A.C. Harley

## Abstract

Glioblastoma is the most common primary malignant brain tumor with a five-year survival rate less than 5%. The standard of care involves surgical resection followed by treatment with the alkylating agent temozolomide (TMZ). GBM cells that evade surgery eventually become resistant to TMZ and lead to recurrence of tumors in patients. With only four drugs currently FDA-approved for GBM treatment, there is a need for a clinically relevant model capable of accelerating the identification of new therapies. Microgels are microscale (∼10–1,000 μm) hydrogel particles that can be used to encapsulate cells in a tailorable 3D matrix. Microdroplets offer short diffusion lengths relative to conventional hydrogel constructs (>1 mm) to limit spatial distributions of hypoxia and potentially screen therapeutics in a controlled and physiologically relevant environment. Here, we establish a method to encapsulate GBM cells in gelatin and polyethylene glycol (PEG) microgels. We show that microgel composition can affect cell morphology and further, that collections of GBM-laden hydrogels can be used to quantify the effect of single vs. metronomic doses of TMZ. GBM metabolic activity is maintained in microgel culture and GBM cells display drug response kinetics similar to previously established literature using macro-scale hydrogel constructs. Finally, we show microgels can be integrated with a liquid handler to enable high-throughput screening using cell-laden microgels.

## 1. Introduction

Glioblastoma (GBM) is the most common and deadly primary malignant brain tumor with a five-year survival rate less than 5% [1, 2]. The current standard of care involves maximal safe surgical resection followed by radiotherapy and adjuvant temozolomide (TMZ) [2–4]. Although treatment increases median survival to 14 months, most patients relapse due to the invasion of therapy-resistant GBM cells beyond the surgical margin [5–8]. A significant challenge for improved clinical outcomes is due to slow progress of therapeutic advancement from lab to clinic [9–11]. Patient-derived xenografts (PDX), where patient tumor cells are implanted into immunocompromised mice, have shown the ability to preserve primary tumor heterogeneity and molecular composition [12–14], these studies can be costly and time-intensive, making them incompatible with many modern high-throughput drug screening (HTS) approaches. Whereas HTS offers the potential to rapidly screen a wide range of doses or drug combinations, the vast majority of these studies rely on two-dimensional (2D) cell culture that cannot recapitulate aspects of the native tumor microenvironment. The GBM tumor microenvironment includes a complex and evolving extracellular matrix (ECM) as well as a range of central nervous system (e.g., pericytes, microglia, astrocytes) and peripheral (e.g. macrophages) cells known to influence disease progression and drug resistance [15, 16]. While hydrogels have been implemented to mimic relevant cell-cell and cell-matrix interactions [17, 18] and are beginning to be used to explore drug resistance [19–22], many current efforts have used either short time frames or supraphysiological compound doses. Thus, there is a critical need for drug screening platforms amenable to HTS approaches that can also replicate aspects of the tumor microenvironment.

Our group has previously developed methacrylamide-functionalized gelatin (GelMA) hydrogels to investigate patterns of GBM invasion and drug response using both conventional cell lines and patient-derived xenograft cells [21, 23–25]. However, the majority of these studies have been conducted in large (∼1mm scale) hydrogel constructs whose fabrication is labor intensive and can require large amounts of both hydrogel material and cell populations. Their large size (1 mm thick; ∼5 mm in diameter) can result in diffusional limitations creating a hypoxic core and hindering cell metabolism [26, 27]. Comparatively, microgels are hydrogels formed as microscale (∼10-1,000 μm) particles that can be used either as distinct units or jammed assemblies referred to as granular hydrogels [28]. Granular hydrogels possess inherent characteristics such as injectability and porosity as well as well-defined rheological properties [29] that have made them an appealing platform for biomedical applications [30]. To date they have been predominantly used as acellular hydrogel particles with cells cultured in voids between particles [31–34]. The encapsulation of cells in microgels has emerged as a versatile tool for tissue engineering and therapy delivery [35–37]. Microgels can be rapidly formed using smaller quantities of starting materials and are not large enough to induce diffusional limitations that reduce the delivery of nutrients, oxygen, or drugs to encapsulated cells [38]. Furthermore, the matrix composition as well as the identity and concentration of cells encapsulated into the microgels can be altered to create discrete microgel populations that can be then mixed [39, 40] to create a more complex model of the tumor microenvironment.

We have recently adapted our gelatin hydrogel system via the inclusion of maleimide crosslinking moieties along the gelatin backbone (GelMAL) to encapsulate primary cells for extended in vitro culture [37]. Here, we evaluate the degree to which hydrogel microdroplets can be integrated with HTS approaches to establish an in vitro GBM drug screening platform. GBM cells are encapsulated in microgels via microfluidic polymerization for precise control over particle size and cell density. We demonstrate microfluidic approaches to monitor cell activity post-encapsulation in vitro as well as successful manipulation (microgel pipettability) via a liquid handler to enable integration with conventional HTS workflows. And finally, we report GBM response to physiologically-relevant doses of TMZ on encapsulated GBM cells.

## 2. Materials and Methods

### 2.1. Gelatin Maleimide (GelMAL) Synthesis and Characterization

GelMAL was synthesized as previously described [37, 41]. Briefly, porcine gelatin type A, 300 bloom (Sigma Aldrich, St. Louis, MO) was dissolved in a mixture of 4:5 (v/v) dimethyl sulfoxide (DMSO; Sigma-Aldrich): aqueous medium of pH 6.0 1M 2-morpholinoethanesulfonic acid buffer (MES; Gold Biotechnology, St. Louis, MO) at 40°C. Then, 6x excess N-succinimidyl 3-maleimidopropionate (Tokyo Chemical Industry Co., Ltd) was dissolved in DMSO and added to the vial containing dissolved gelatin and capped. The pH of the reaction was then adjusted to 4.5 using 1.0 N HCl. Reaction proceeded for 24 h at 40°C 720 RPM, the solution was dialyzed against distilled water acidified to pH 3.25 using 1.0 N HCl, at 40 °C for 5 days. Product was frozen lyophilized, then stored at -20 °C until further use. ^1^H NMR was used to measure the degree of functionalization (DOF). GelMAL with DOF between 49 and 51% (data not shown) was used in this study.

### 2.2. Flow-focusing Microfluidic Device Construction

Polydimethylsiloxane (PDMS) flow-focusing microfluidic devices with a 200 μm nozzle were cast from either silicon or SU8 masters using the Sylgard 184 silicone elastomer kit (Dow Corning, Midland, Michigan) [36, 37]. Ports were incised using a 1 mm biopsy before being bonded directly to glass slides after plasma treatment.

### 2.3 U87MG Cell Culture

U87MG cells (American Type Culture Collection [ATCC], HTB-14) were maintained in DMEM, high glucose (Thermo Fisher Scientific, Waltham, MA) supplemented with 10% (v/v) fetal bovine serum (FBS; R&D Systems, Minneapolis, MN) and 1% (v/v) antibiotic-antimycotic (Thermo Fisher Scientific) in a humidified 5% CO_2_ incubator at 37°C.

### 2.4 Cell Laden Microgel Fabrication

GelMAL microgels were formed using a flow-focusing microfluidic device with a 200 μm nozzle [36, 37]. 3 vol% SPAN80 (Sigma-Aldrich) in light mineral oil (Sigma-Aldrich) was used as an oil mixture. 1,4-dithiothreitol (DTT) (Sigma-Aldrich) was freshly prepared at 20 mg/mL aqueous solution emulsified with the oil mixture at a ratio of 1:15 (v/v). 4.8 wt% GelMAL was dissolved in PBS with 12% (v/v) OptiPrep (Sigma Aldrich) and 0.06% (w/v) Pluronic F-108 (Sigma Aldrich). Addition of OptiPrep and Pluronic F-108 was incorporated to increase the cell medium density and aid cell distribution. U87MG cells were passaged using TrypLE Express (Thermo Fisher Scientific) before treating with ethylenediaminetetraacetic acid (0.52 µM) (EDTA; Thermo Fisher Cat# 15575020) for 15 minutes at 37°C. Cells were washed with PBS three times and kept on ice until mixed with the GelMAL precursor. The GelMAL precursor and cells were mixed at a 1:5 ratio (final cell dilution: 2 x 10^6^ cells/ mL), respectively. The oil mixture, DTT crosslinker, and gel precursor were loaded into syringes then independently pumped into the microfluidic device using syringe pumps (Pump 11 Elite, Harvard Apparatus, Holliston, MA) at rates of 0.5 μL/ min (oil mixture), 30 μL/ min (crosslinker emulsion), and 5 μL/ min (gel precursor + cells), respectively. When encapsulating cells, the volume of precursor pumped through the device was limited to 60 μL to limit the total time cells were exposed to shear stress from the pumps. Microgels were collected in culture medium on an ice bath then shaken at 60 rpm at 4°C for 20 minutes to allow completion of crosslinking. Microgels were then washed once with PBS, 0.1% (v/v) Tween-20 (Fisher Scientific) in PBS, and culture medium. Washes were performed at 300 rcf x 3 minutes of centrifugation. PEG microgels were formed as stated previously utilizing PEG-4MAL (JenKem Technology USA Inc., Plano, TX). PEG hydrogels were created with the inclusion of 0.3 mM RGD peptide (MedChem Express, GRGDSPC) containing a free cysteine to facilitate crosslinking with the PEG-4MAL monomer (added before mixing with cells).

### 2.5 Viability Measurement of Encapsulated Cells

Non-encapsulated cells and cell-laden microgels were stained with live/dead indicators calcein AM and BOBO-3 iodide (Thermo Fisher Scientific, R37601) and Hoechst 33342 (1:1000) for 15 min before washing and imaging. Microgels were imaged using a DMi8 Yokogawa spinning disk confocal microscope equipped with a Hamamatsu EM-CCD digital camera (Leica Microsystems, Inc., Deerfield, IL).

### 2.6 Measurement of DNA Content

Double-stranded DNA (dsDNA) was extracted from non-encapsulated cells and cell-laden microgels using the QIAamp DNA Micro Kit (QIAGEN, Hilden, Germany) following the manufacturer’s protocol, with the following modifications: resulting lysate was passed through a QIAshredder (QIAGEN). The lysate was then transferred to the QIAamp column, and DNA was extracted according to the kit instructions and eluted in molecular-grade water. Quantification of dsDNA was performed using the Quant-iT PicoGreen dsDNA Assay Kit (Thermo Fisher Scientific), and fluorescence was measured using a Synergy LX Multi-Mode Microplate Reader (BioTek, Winooski, VT) at 485 nm excitation wavelength and 528 nm emission wavelength.

### 2.7 Tracing of Morphological Changes in GelMAL Microgels

Microgels were injected into a microfluidic chip (idenTx 3, AIM Biotech, Singapore) to maintain in place and trace morphological changes. Culture medium was changed daily following the manufacturer’s protocol. Microfluidic chips were placed in a humidity chamber in an incubator at 37°C. Microgels were imaged using a DMi8 Yokogawa spinning disk confocal microscope equipped with a Hamamatsu EM-CCD digital camera (Leica Microsystems).

### 2.8 Tracking of Metabolic Activity of Cell-laden GelMAL Microgels

Cell-laden GelMAL microgels were maintained under the previously described culture conditions. Metabolic activity was assessed using alamarBlue Cell Viability Reagent (Thermo Fisher Scientific). The reagent was added to each well to a final concentration of 10% (v/v) and incubated for 30 minutes in the dark at 37 °C with 5% CO_₂_. Fluorescence was measured using a Synergy LX Multi-Mode Microplate Reader (BioTek) at an excitation wavelength of 530 nm and an emission wavelength of 590 nm. Blank controls, including wells containing acellular microgels and wells with only media, were included for each treatment condition.

### 2.9 Proliferation Analysis of Encapsulated Cells

The fraction of proliferating cells was assessed using the Click-iT EdU Cell Proliferation Kit for Imaging (Thermo Fisher Scientific) to measure the fraction of EdU+ cells in microgel culture. On day 2, 3, and 4, a media change was performed for each sample and half the media was replenished with fresh culture medium. Other half of medium was introduced to make a final concentration of 10 µM EdU. Microgels were incubated with EdU for 24 h, after the media was removed and microgels were fixed in formalin for 15 min. The click reaction to attach Alexa Fluor 647 to the alkyne-containing EdU was performed per manufacturer’s instructions. Microgels were stained with Hoechst 33342 (1:1000) for 15 min before washing and imaging.

### 2.10 TMZ Drug Response

Cell-laden microgels were cultured in suspension for three days in complete media. After three days, microgels were manually pipetted into a 96 well plate with each well containing approximately 4.5 x 10^4^ cells. Temozolomide (TMZ, Selleck Chemicals, Houston, TX) dissolved in 130 mM DMSO (Sigma Aldrich) was diluted in culture medium and added to wells. 0.0077% v/v DMSO was used as a vehicle control. Changes in cell activity were assessed using alamarBlue Cell Viability Reagent (Thermo Fisher Scientific). For GelMAL microgels, a single dose of TMZ was given on day three and changes in cell behavior were assessed two days post-treatment. PEG-4MAL microgels were placed into an HTS Transwell-96 permeable support plate (Corning, Tewksbury, MA) to enable treatment supplements and ease medium changes. PEG-4MAL microgels were treated for five days consecutively beginning on day three. Changes in cellular activity were assessed on day five, seven, and nine post-encapsulation.

### 2.11 Morphological Changes in Microgel Culture

Morphological changes in the microgel cultures were assessed by staining actin filaments using phalloidin. Cell-laden microgels were cultured in free suspension then fixed in formalin for 15 minutes at room temperature on the respective day. Following fixation, samples were permeabilized with 0.5% Triton X-100 in PBS for 15 minutes. Phalloidin conjugated to Alexa Fluor 488 (1X final concentration from a 400X stock in DMSO; Thermo Fisher Scientific) was diluted in PBS and incubated with the microgels for 30 minutes at room temperature in the dark. Microgels were stained with Hoechst 33342 (1:1000) for 15 min before washing and imaging.

### 2.12 Liquid Handler Movement of Microgels

Cell-laden PEG microgels were cultured for three days in suspension. On day three, microgels were pipetted into a 384 well plate using the Biomek i5 automated liquid handler (Beckman Coulter Life Sciences, Indianapolis, IN) with each well containing approximately 4.5 x 10^4^ cells using facilities within our campus High Throughput Screening Facility (Cancer Center at Illinois). Cell-laden PEG-4MAL microgels were manually pipetted into a 384 well plate as a control. Cell activity was measured at two days post-transfer using alamarBlue Cell Viability Reagent (Thermo Fisher Scientific). Microgel integrity was measured by imaging microgels automatically moved by a liquid handler or manually with a micropipette using the EVOS M5000 Imaging System (ThermoFisher Scientific).

### 2.13 Statistics

Statistics were performed using GraphPad Prism (GraphPad Software, Inc., Boston, MA). %LIVE population, double stranded DNA levels, metabolic activity, and EdU+ population are reported as the mean ± standard error of the mean. Normality of data was determined using the Shapiro-Wilk test, and equality of variance was determined using Brown-Forsythe Test. Data satisfying normality was analyzed using the unpaired t-test or one-way analysis of variance (ANOVA) followed by Tukey’s post hoc test. When data were not normal, comparisons between multiple groups were performed using a Kruskal-Wallis test with Dunn’s post-hoc. Significance was determined as p < 0.05. A minimum of three biological samples and three technical replicates was used for all experiments.

## 3. Results

### 3.1. A method to reliably encapsulate GBM cells in GelMAL microgels and retain viability post-encapsulation

We established a consistent and robust method to rapidly encapsulate GBM cells into nanoliter-volume hydrogel microdroplets **(Fig 1A)**. U87MG GBM cells were mixed (2 x 10^6^ cell/ mL) with a 4% GelMAL precursor solution then run through a flow-focusing microfluidic device; this cell concentration was chosen based on prior work [39] to result in 2 – 5 cells per microgel to provide an environment for cell proliferation and to assess therapeutic efficacy. Microgels were collected and washed then cultured in suspension for processing. Consistent with prior literature [42], we observed maintaining cell viability after microfluidic emulsion required limiting the length of each fabrication run to reduce the level of shear stress experiences by traveling to and through the microfluidic device. For runs of up to 12 minutes (60 μL of cell and precursor suspension pumped at a rate of 5 μL/ min), U87MG cell viability displayed an immediate ∼20% decrease **(Fig 1B).** Although encapsulation process induced an immediate loss of viability, the live fraction remains stable at 1- and 3-days post-encapsulation **(Fig 1C)**, suggesting the initial cell loss does not affect the long-term culture of our microgels.

**Figure 1.**
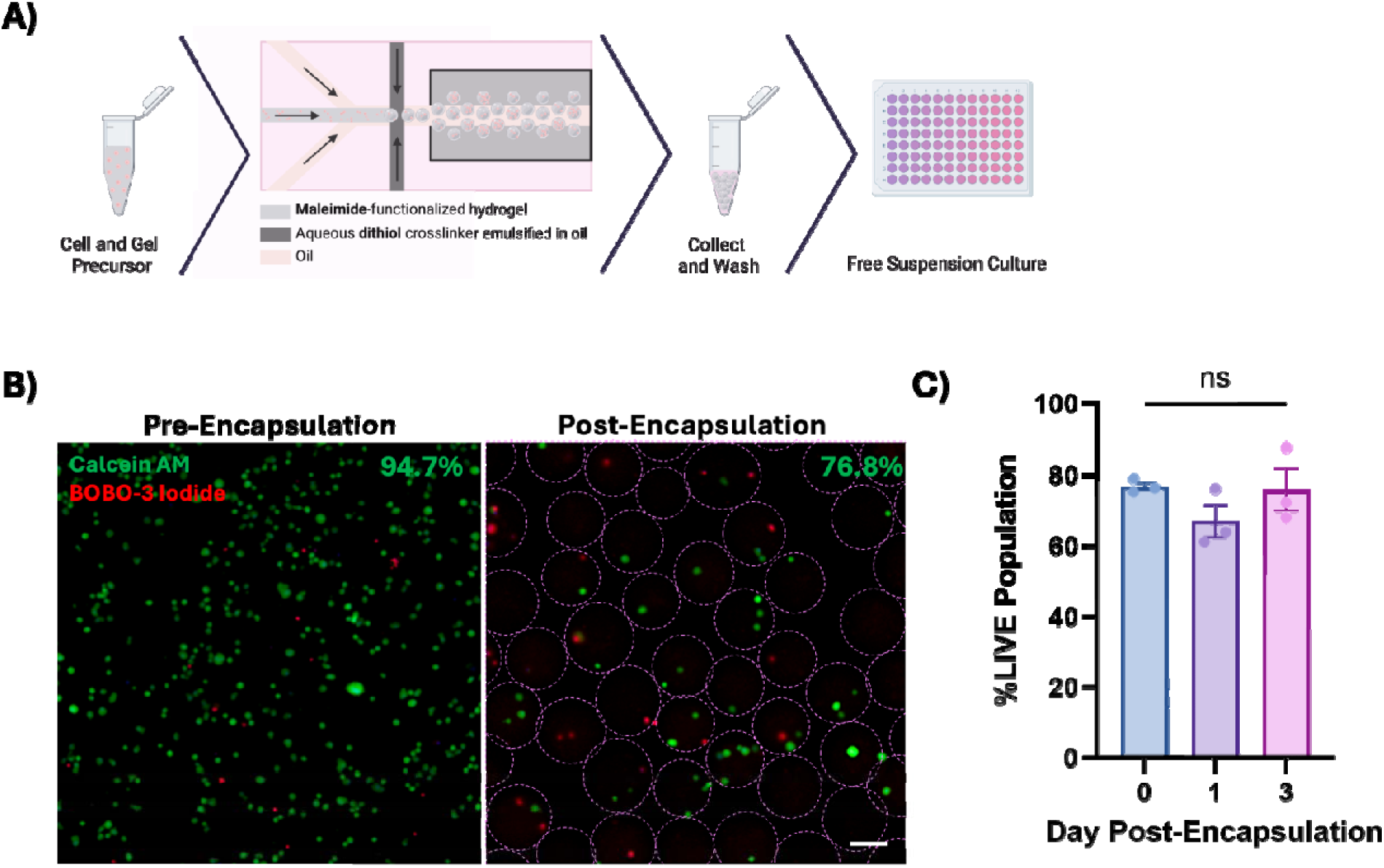
**A)** Method schematic of cellular encapsulation. Cells are mixed with the polymer precursor then run through a microfluidic device. Microgels are collected, washed, and cultured in free suspension. **B)** Viability of cells before encapsulation (94.7%) and directly after encapsulation (76.8%). **C)** Viability on days 0, 1, and 3 post-encapsulation (76.8, 67.0, and 76.1%, respectively). Scale bar= 100 μm.

### 3.2. Characterization of cellular behavior post-encapsulation

Next, we set out to define how cellular behavior is impacted by the encapsulation process. We assessed morphological changes after encapsulation, using cell-laded droplets packed into an idenTx 3 microfluidic chip to make repeated measurements of GBM cells **(Fig 2A)**. On the initial day of encapsulation (Day 0), encapsulated cells were largely spherical in shape consistent with isolation and microfluidic encapsulation. On day 1, we observed cells begin to display protrusions which become more pronounced by day 3. This elongated morphology is comparable to the morphology observed in both 2D cell culture as well as 3D gelatin hydrogel culture for the U87MG cell line.

**Figure 2.**
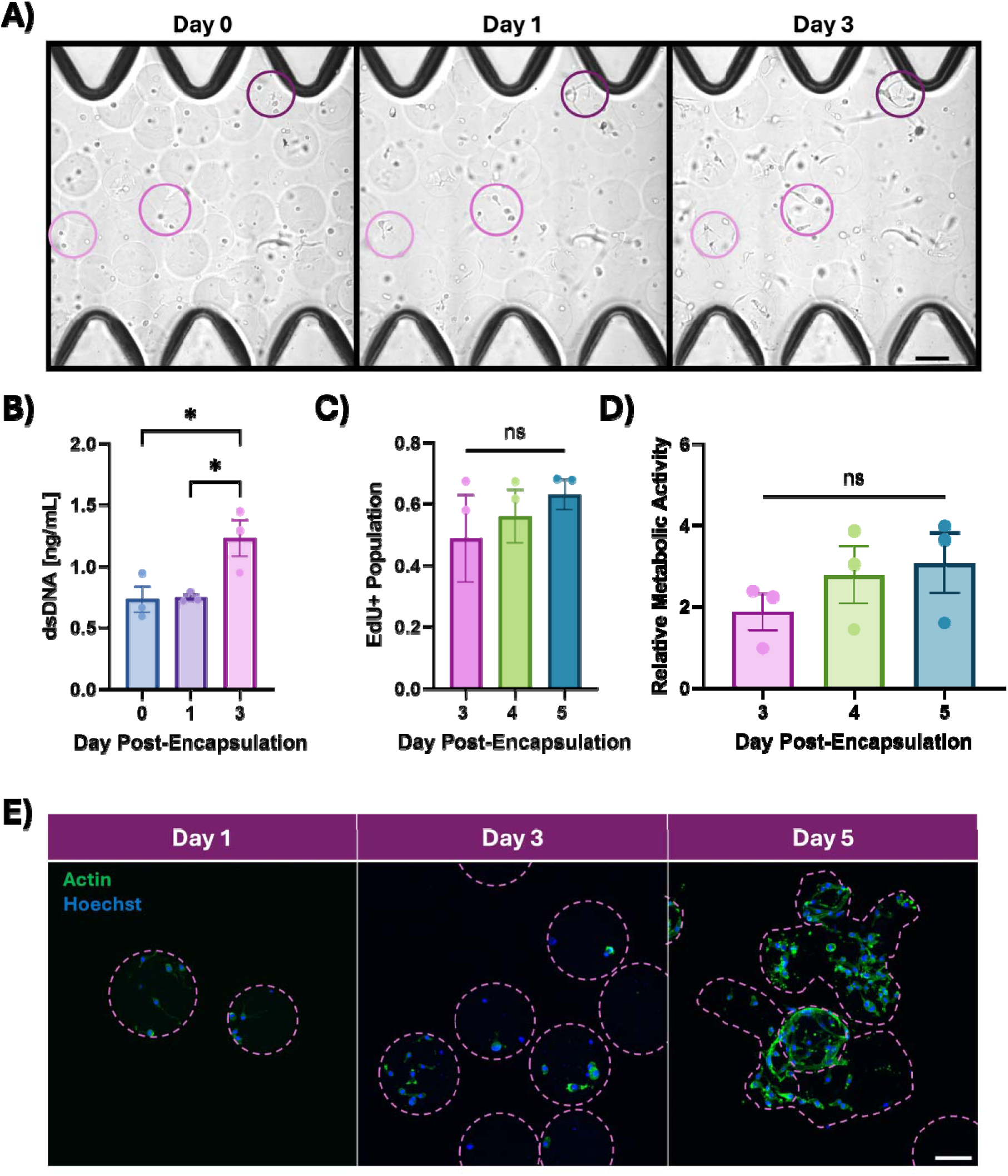
**A)** Brightfield images of GelMAL microgels in idenTx 3 microfluidic chip at days 0, 1, and 3. **B)** Double-stranded DNA (dsDNA) levels in microgels at days 0, 1, and 3 post-encapsulation (0.73, 0.75, and 1.3 ng/ mL, respectively). Microgel levels of **C)** 5-ethynyl-2’ – deoxyuridine (EdU+) incorporated (0.49, 0.56, and 0.63%, respectively) and **D)** levels of cellular activity (1.86, 2.78, 3.07-fold change, respectively) at days 3, 4, and 5 post-encapsulation. **E)** Actin stain demonstrating cellular morphology in GelMAL microgels at day 1, 3, and 5. Scale bar= 100 μm.

We also investigated intracellular activities to further understand the cellular behavior inside microgel culture. Double stranded DNA (dsDNA) levels suggest cell proliferation begins on day 3 where we see a significant increase of dsDNA compared to the initial population on day 0 **(Fig 2B)**. Past day 3, we see a trend of increasing proliferating (Edu+) population and metabolic activity (**Fig 2C and 2D**, respectively). Actin staining of U87MG cells in GelMAL microgels cultured in free suspension **(Fig 2E)** reveals an increase in cell numbers and formation of cell aggregates, as well as a trend that suggests (by day 5) GBM cell growth may lead to significant GelMAL microgel remodeling and merging. This suggests that analysis of GBM cell culture in GelMAL microgels may have a maximum timepoint of 5 days.

### 3.3. Assessing single-dose drug response in GelMAL microgel culture

Based on our work that showed GelMAL microgels supported 5-day cell culture, we then performed single TMZ dose response assays to establish a microgel drug screening platform **(Fig 3A)**. Microgels were cultured for 3 days to allow time for GBM cell proliferation to initiate before dosing microgel in culture with TMZ [0, 10, 100 µM]. Drug efficacy was assessed by changes in cell metabolism 2-days post-treatment. U87MG cells encapsulated in GelMAL microgels did not show a change in activity when exposed to a low (sub-physiological) dose of TMZ [10 µM] **(Fig 3B)**. However, cells exposed to a higher dose of TMZ [100 µM] showed a significant decrease in metabolic activity **(Fig 3B)**.

**Figure 3.**
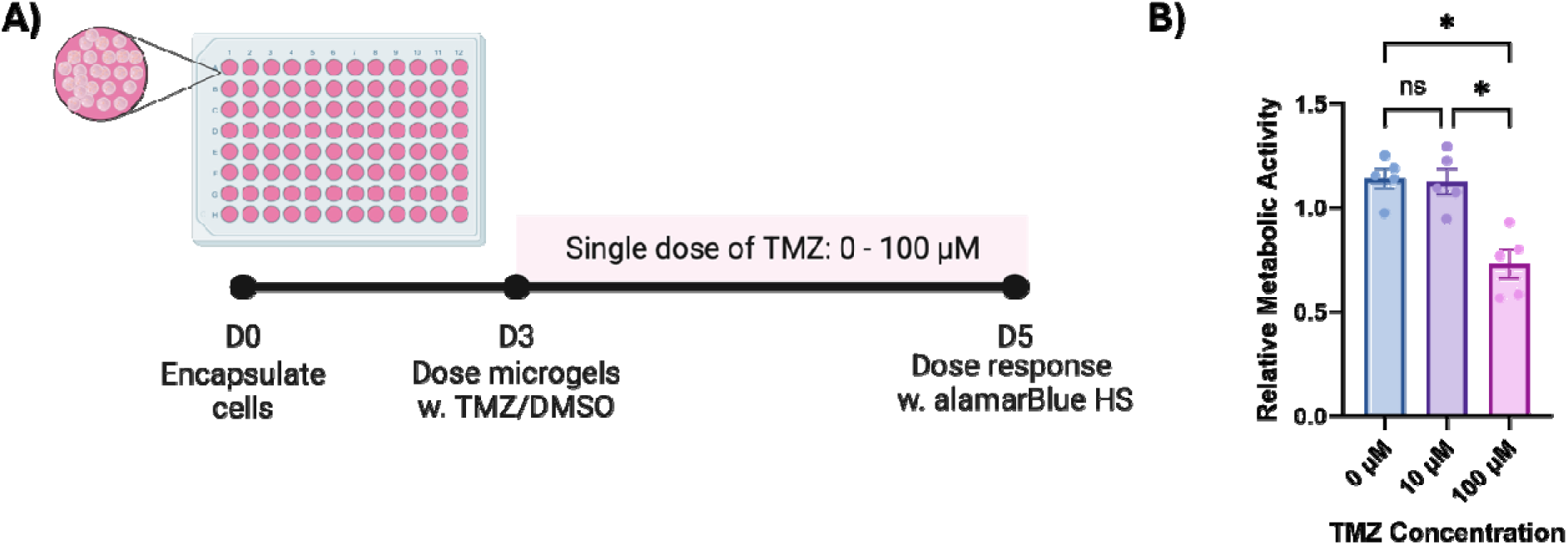
**A)** Timeline for single dose response of GelMAL microgels. Microgels are cultured for three days then a single TMZ dose is exhibited to cells and response is recorded two days post-treatment. **B)** Cellular activity of microgels two days post-TMZ treatment (1.14, 1.13, and 0.73 fold-change, respectively).

### 3.4. Enabling metronomic dosing in PEG microgel culture

To overcome challenges of significant cell-mediated remodeling of GelMAL microdroplets, we also encapsulated U87MG GBM cells in PEG-4MAL microdroplets. U87MG cells were encapsulated in RGD-functionalized PEG-4MAL, using the same process as GelMAL hydrogels, then cultured for up to 9 days post-encapsulation. Actin staining of GBM cells in PEG microgels revealed GBM cells form multicellular aggregates within microgels that increased in size with time **(Fig 4A)**, likely a consequence of the lack of the non-degradable nature of PEG-4MAL microgels that would otherwise keep cells constrained. We used the extended culture time of PEG-4MAL microgels to examine the effect of metronomic daily dosing of TMZ **(Fig 4B)**. GBM-laden PEG microgels were cultured for 3 days after encapsulation in conventional culture, then were given either a single dose of 100 µM TMZ on day 3 or five daily consecutives doses of 20 µM TMZ (days 3-7). We assessed changes in cell metabolic activity on day 5, 7, and 9 (2, 4, and 6 days after the initial TMZ dose was administered). We observed trends towards reduced cell metabolic activity as a result of metronomic dosing at days 5 and 7 and a significant reduction in GBM cell metabolic activity on day 9 compared to DMSO control or a single supraphysiological dose of TMZ **(Fig 4C, D, E)**.

**Figure 4.**
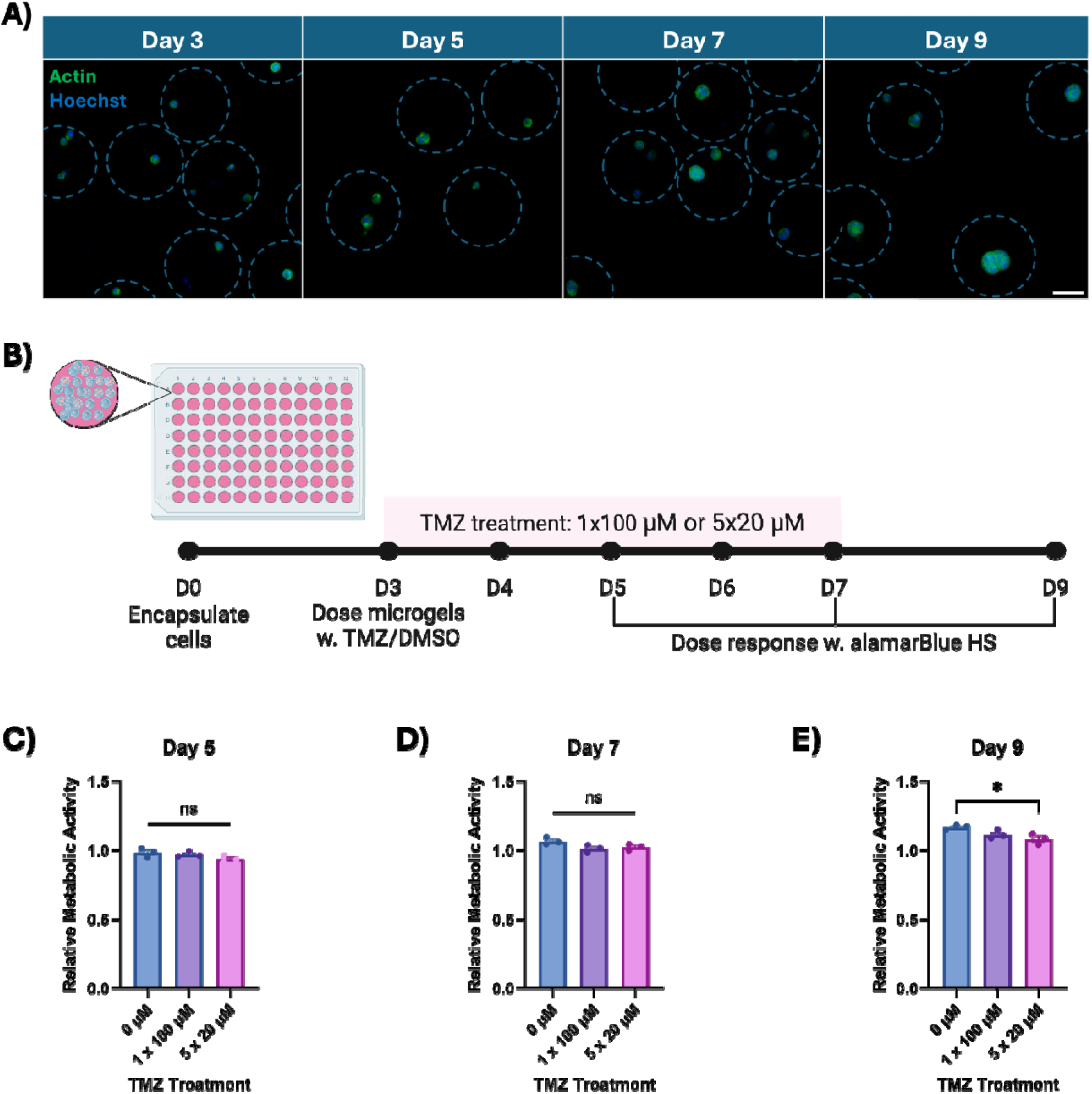
**A)** Actin stain demonstrating cellular morphology in PEG-4MAL microgels at day 3, 5, 7, and 9. **B)** Timeline for metronomic dose response of PEG-4MAL microgels. Microgels are cultured for three days then treated with a single dose of 100 μM TMZ or five consecutive daily doses of 20 μM TMZ and response is recorded two days post-treatment. Cellular activity of microgels on **C)** day 5 (0.98, 0.97, and 0.93 fold-change, respectively), **D)** day 7 (1.06, 1.01, and 1.02 fold-change, respectively), and **E)** day 9 (1.17, 1.11, and 1.08, respectively). Scale bar=100 μm.

### 3.5. Evaluating the handleability of microgels with a liquid handler to explore high-throughput screening

Successful extended culture and the ability to measure the therapeutic efficacy of alkylating agents (e.g., TMZ) at physiological doses suggest the microgel system may have significant potential to automate testing of large compound libraries against cells within highly controlled and physiologically relevant tissue microenvironments. U87MG cells were encapsulated in PEG-4MAL microgels, allowed to culture in suspension for 3 days, then 10 µL of PEG-4MAL microgels was moved automatically by a liquid handler or manually with a micropipette into a 384 well plate containing 20 µL of media per well **(Fig 5A)**. We first assessed the integrity and structure of microgels post-movement, finding microgel shape and cellular integrity was maintained after liquid handler and micropipette **(Fig 5B)**. Microgels were then maintained in 384 well plate culture for two days before we assessed cell metabolic activity, finding that microgels that were moved with the liquid handler displayed significantly metabolic activity compared to those moved manually via micropipette **(Fig 5C)**. These data confirm the ability to integrate GBM-laden microgels with high throughput screening (HTS) automation to expand the scope of drug testing in physiologically-relevant matrix environments.

**Figure 5.**
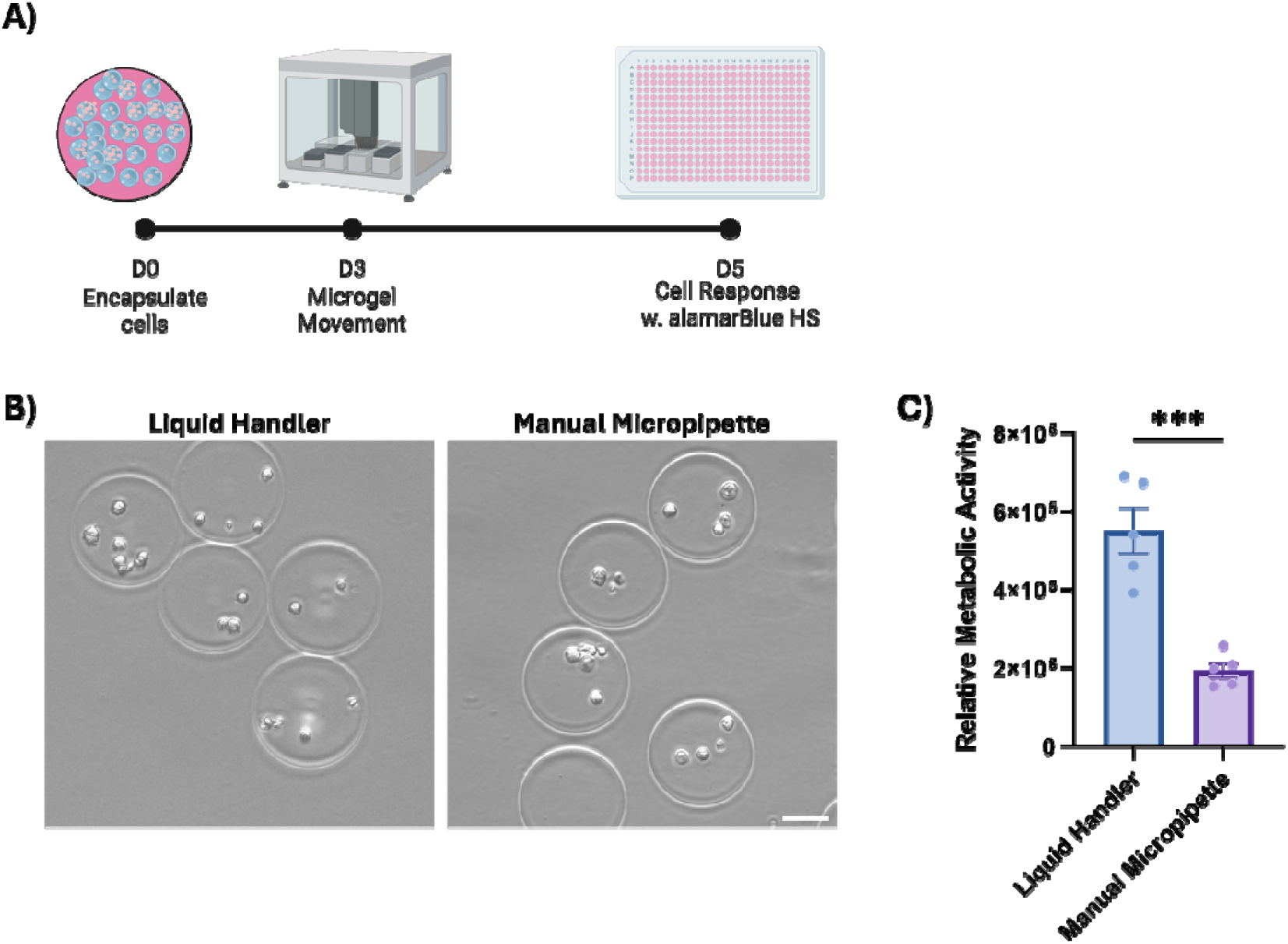
**A)** Timeline for liquid handler experiments. Microgels are cultured for three days before movement either by liquid handler or manual pipette. Cellular response to movement is recorded two days post-movement. **B)** Microgel structure post-movement manually by micropipette or automatically by liquid handler. **C)** Metabolic activity of microgels two days post-movement by either manual micropipette or liquid handler (5.5 x 10^5^ and 1.9 x 10^5^ fold-change, respectively). Scale bar= 100 μm.

## 4. Discussion

A significant challenge to improving survival rates for glioblastoma remains the identification of new therapeutic approaches. While TMZ improves median survival to ∼14 months [6], however, it remains the most deadly primary malignant brain tumor. Reliable and predictive preclinical models that recapitulate aspects of the local microenvironment offer the potential to accelerate the discovery of GBM treatments. Hydrogels are highly tunable and can be tailored to study cell-cell and cell-matrix interactions in a highly controlled environment. Previous studies have encapsulated GBM cells in bulk hydrogels to measure the impact of biomaterial stiffness [43–45], structure [46, 47], and composition [48–50] on morphology, migration, GSC maintenance, and drug response [51–54]. However, due to their size and the relatively complex process of making and moving large hydrogel volumes between culture conditions, macro-scale hydrogels can exhibit biotransport limitations [55] and may not accommodate small cell populations often obtained from patient specimens. The encapsulation of cells into microgels offers an appealing alternative to overcome potential biotransport limitations typical in large hydrogel networks [56] and to provide a pathway to use cell-laden microgels as a building block for high throughput drug screening.

We established a robust and consistent method to encapsulate GBM cells into nanoliter-volume GelMAL and PEG-4MAL microdroplets. Microgel fabrication utilizing microfluidic polymerization allows for precise control over particle size and shape compared to other methods [35, 36, 57]. However, achieving high cell viability within microfluidic emulsion is limited by the level of shear stress cells are exhibited to while traveling to and through the microfluidic device [42]. By limiting fabrication runs to <15 minutes we describe an encapsulation method that retains approximately 80% of the live population that exhibits subsequent growth (dsDNA, metabolic activity) in culture. Interestingly, with cellular health intact and a growing demand for culture area, GBM cells showed significant remodeling associated changes to GelMAL microgels, leading to the formation of interconnected cell-microgel constructs by day five of culture. These results are consistent with macroscale studies of GBM-hydrogel interactions, where GBM cells showed significant matrix remodeling capacity [27, 58, 59]. While potentially interesting from the perspective of using initial microgel cultures as a granular niche to support de novo generation of interconnected tissue constructs at a much larger scale, we chose to limit Gel-MAL microgel culture for this work to the period where the biomaterial environment remained cell-laden GelMAL microgels. While RGD-functionalized PEG-4MAL microgels enabled extended culture of GBM cells, GBM cells in PEG-4MAL microgels formed multi-cellular aggregates. Consistent with a PEG network that does not contain enzymatically degradable motifs but does contain RGD cell adhesion sites, we did not observe evidence of microgel degradation but rather cell aggregation within the hydrogel matrix. And while not shown, we observed in cases of extended (>12 days), GBM cell laden microgels exhibited rupture, suggesting extended cell proliferation without remodeling fractured the microgel network. Future extensions of this work could explore the use of PEG or hyaluronic acid functionalized PEG microgels containing enzymatically degradable motifs in order to explore cell mediated remodeling processes. While such materials may be of value for studies of GBM invasion, the viability and proliferative activity of GBM cells in PEG-4MAL and GelMAL microgels made them an ideal substrate to examine responses to physiologically relevant therapeutic strategies.

Temozolomide (TMZ) is an alkylating agent used to treat GBM [60, 61]. While delivered as a pro-drug, it undergoes hydrolysis that induces DNA methylation and induce cellular apoptosis. The methylation of DNA is thought to be the principal mechanism responsible for TMZ cytotoxicity [60–63]. TMZ acts on proliferative cells, which has led many in vitro studies to rely of supraphysiological TMZ doses to induce rapid changes in cell viability amenable to conventional in vitro analyses. Recently, we used extended (>7 day) in vitro culture to measure response to physiologically-relevant (<30 µM) and metronomic TMZ doses in GelMA macrogels [64].

Adapting this protocol, here we encapsulated U87MG GBM cells in GelMAL or PEG-4MAL microgels then cultured the cell-laden microgels for three days before TMZ treatments. In GelMAL microgels, U87MGs were unaffected after 48 hours at a low, but physiologically relevant dose of 10 μM TMZ, but showed significant reduction in metabolic activity 48 hours after a larger dose of 100 μM TMZ. These results are consistent with prior findings for U87MG in 2D culture that showed the half maximal inhibitory concentration (IC50) of TMZ ranges from 92.0 to 590.1 μM [65]. Hence, we demonstrate U87MG cells in microgel culture reveal similar trends in response to low IC50 doses of TMZ.

While measuring dose-response to supraphysiological TMZ dosages is an important first finding, TMZ concentrations in patient samples often register below 10 μM [66–68]. To achieve the antitumor effect of TMZ, patients instead receive metronomic administrations (e.g., TMZ orally 5 days per week for 4 weeks) [69, 70]. We previously showed macroscale gelatin hydrogels enabled the study of metronomic TMZ dosages [64], but the GelMAL microgel degradation limited the length of our studies. We used RGD-functionalized PEG microgels to extend our experimental timeline to accommodate microgel pre-culture, 5 days of metronomic dosing, then 2 additional days of culture to register responses. Cells were cultured for three days before being treated by either a single 100 μM dose or five consecutive daily doses of 20 μM for a cumulative dose 100 μM TMZ. Interestingly, metronomic dosing exhibited a cumulative negative effect on GBM metabolic activity while a single 100μM dose did not. While different than the significant effect of 100 μM TMZ on U87MGs in GelMAL microgels, the change in microgel chemistry is likely inducing changes in U87MG cell activity. Indeed the 100 μM dose is still at the low end of reported IC50 doses in 2D culture (92.0 to 590.1 μM [65]), and the efficacy of a single TMZ dose requires significant levels of cell proliferation at the moment of treatment. Here, the ability to pursue extended study of the effect of metronomic dosing is made possible by cell-laden PEG microgels, offering a platform for future studies using physiologically-relevant drug dosages. Here, future studies can again take advantage to advances in matrix engineering via the use of PEG-4MAL/GelMAL mixtures or the inclusion of additional component of the tumor extracellular microenvironment such as hyaluronic acid [49, 71, 72] to create an environment tailored for cell remodeling, proliferation, and drug testing.

The success of PEG-4MAL microgels for examining metronomic TMZ dosing inspired efforts to integrate microgels with infrastructure essential for high-throughput compound screening. Here, GBM cells encapsulated in hydrogel microgels could be manipulated via a liquid handler into well-plate formats. GBM-laden PEG-4MAL microgels were cultured three days prior to liquid handler manipulation to assess changes in cellular activity vs. (low-throughput) manual pipetting typically used for microgel cultures. Microgel integrity was retained after liquid handler manipulation with U87MG microgels showing a higher cellular activity after liquid handler (vs. manual pipetting) manipulation. While intriguing, this result may be linked to consistency in handling, with the liquid handler efficiently pipetting the microgels at a consistent rate (uptake: 5 μL/sec; dispense: 20 μL/sec) rather than the via manual processes, resulting in batch to batch consistency. These results provide confidence for future use of PEG-4MAL microgels for high capacity and efficient studies of drug discovery using cells maintained in well-defined 3D matrix environments.

## 5. Conclusions

Tissue engineering models of the tumor microenvironment have provided a valuable tool to investigate processes of GBM cell matrix remodeling, invasion, and drug response. However, the large size of conventional hydrogel materials makes them not viable for high-throughput compound screening workflows. Here, we describe an approach to encapsulate GBM cells in GelMAL and PEG-4MAL microgels. Microgels culture reveals an increase in cell activity and matrix remodeling with time. GelMAL microgels can be used for single dose response studies using TMZ (<5 days). PEG-4MAL microgels allowed for extended culture and study of clinically relevant metronomic dosing strategies for cells in 3D matrix environments. Lastly, these microgels can be integrated into conventional workflow (automated liquid handler) necessary to use microgels in high-throughput drug screening facilities. Together, this suggests cell-laden microgels offer a valuable system for high-dimensional studies of drug efficacy with cells maintained in well-defined matrix microenvironments.

## Acknowledgements

We acknowledge the following institutes for access to their facilities and services: the Roy J. Carver Biotechnology Center at UIUC, the School of Chemical Sciences High-Throughput Screening Facility, the Tumor Engineering and Phenotyping Core at the Cancer Center at Illinois, the Carle R Woese Institute for Genomic Biology, and Materials Research Laboratory. Research reported in this publication was supported by the National Cancer Institute of the National Institutes of Health under Award Number R01 CA256481 (BACH), the National Institute of Diabetes and Digestive and Kidney Diseases of the National Institutes of Health under Award Number 2 R01 DK099528 (BACH), as well as the National Institute of Biomedical Imaging and Bioengineering of the National Institutes of Health under Award Number T32EB019944. The content is solely the responsibility of the authors and does not necessarily represent the official views of the NIH. The authors are also grateful for additional funding provided by the Department of Chemical & Biomolecular Engineering, the Carl R. Woese Institute for Genomic Biology, the Cancer Center at Illinois, and the Illinois Scholars Undergraduate Research Program at the University of Illinois Urbana-Champaign.

## Contributions (CRediT: Contributor Roles Taxonomy [73, 74])

**B. A. Payan:** Conceptualization, Data Curation, Formal Analysis, Visualization, Investigation, Methodology, Writing – original draft, Writing – review & editing.

**A. Carrillo Diaz De Leon:** Data Curation, Investigation, Methodology, Writing – original draft

**T. Anand:** Visualization, Investigation, Methodology

**M. Mora-Boza:** Methodology

**V.V. Krishnamurthy:** Methodology

**G.B. Thompson:** Investigation, Methodology

**A.J. García:** Resources, Writing – review & editing

**B.A.C. Harley:** Conceptualization, Resources, Project administration, Funding acquisition, Supervision, Writing – review & editing.

## Notes

### Competing Interest Statement

The authors have declared no competing interest.

